# Individual human genomes frequently contain variants that have evolutionary couplings

**DOI:** 10.1101/2020.08.07.240887

**Authors:** Henry J Martell, Darren K Griffin, Mark N Wass

**Affiliations:** School of Biosciences, University of Kent, Canterbury, Kent, CT2 7NJ, UK

## Abstract

Coevolution has been widely studied between species and has an important role in our understanding of biological function. For proteins there has recently been interest in the identification of positions within proteins that have coevolved including their use for modelling protein structure. Such studies focus on the identification of coevolving positions (or evolutionary couplings) within multiple sequence alignments of proteins from many species. Here, we exploit large human genome resources to investigate if it is possible to use genetic variation data from a single species, human, to identify positions within proteins that have coevolved. We combine the 1000 genome project genetic variation data with protein structural data to identify variant-variant interactions within individual human genomes. We find >4,000 combinations of variants that are located close in 3D protein structure and >1,200 in protein-protein interfaces. Many variant combinations include compensatory amino acid changes (e.g. maintaining charge/functional groups), thus supporting that these are coevolutionary events. Our study highlights that it is possible to identify coevolution within a single species. Given the important role that genetic variation plays in causing disease it is important for variant interpretation and precision medicine to consider the gestalt effects of variants rather than individual variants in isolation.

## INTRODUCTION

In recent years, advances in sequencing technologies have enabled extensive human genetic variation data sets to be generated [1-8]. These resources are extremely rich in evolutionary information and we are far from having an extensive understanding of the genetic variation present within the human genome. One area of interest is the understanding of coevolution to identify positions with the encoded proteins that are evolutionary coupled and therefore likely to have an important role in maintaining protein structure and/or function [9].

Population studies are also beginning to demonstrate how coevolution has occurred during human evolution as demonstrated by “genetic buffering” or “genetic compensation”. Studies have identified individuals carrying variants for severe childhood Mendelian diseases who have reached adulthood with no clinical manifestation of the disease [10], while other studies have shown that variants thought to cause Mendelian diseases are far more common in human populations than would be expected based on disease prevalence [11-15]. This suggests that some individuals with these variants are protected from disease by other variants compensating for the variant that would normally cause disease, representing examples of evolutionary couplings between variants.

This context dependence has long been observed between species, as Compensated Pathogenic Deviations (CPDs) [16], where missense variants that cause disease in humans are wild type in other species. For example, the variant p.Val20Glu in human β-haemoglobin (rs33918474) causes erythrocytosis, whereas p.Glu20 is wild type in horse β-haemoglobin. A second difference at residue 69 is likely to compensate for this, with the horse sequence containing His and the human Gly [17]. In horse β-haemoglobin Glu20 and His69 form a hydrogen bond, while Val20 and Gly69 have van der Waals interactions

Further, it has been estimated that between 3-12% of disease associated variants in ClinVar [8] are observed in orthologs in vertebrates, thus representing possible CPDs [18]. Experimental analysis of CPDs in the genes BBS4, RPGRIP1L, and BTG2 showed that CPDs were able to rescue pathogenic variants and restore wild type function [18]. Another study identified mutually compensatory variants in human p53 that preserve protein stability [19]. In these examples, there are combinations of variants where the destabilising or damaging effects of individual variants are non-additive and the full variant combination better preserves the properties of the wild type protein and provides examples of potential evolutionary couplings.

The identification of evolutionary couplings has to date focussed on comparisons between many different species and in recent years there have been significant advances in their identification and use for the modelling of protein structure and protein-protein complexes [20-31]. Here, we exploit large human genomic datasets to use the variation present within a single species, human, to identify evolutionary couplings. We quantify the extent of possible evolutionary couplings within protein coding regions of the human genome. The chromosomally-phased genomic variation data present in the 1,000 Genomes Project are utilised, generating individual variant sets for each of the 2,504 genomes. We analyse the extent of co-occurring variants within individual proteins, how co-occurring variants are distributed within both protein 3-dimensional structures, and at protein-protein interface sites. Thus, we do the opposite of methods using evolutionary couplings to model protein structure, instead we identify potential evolutionary couplings and use their co-localisation in protein structure or in protein-protein interfaces as evidence that they are evolutionary couplings.

## MATERIAL AND METHODS

### Protein Sequence Data

The UniProt human proteome set (UP000005640) was used, version last modified on 22/11/2015 [32], and for human Ensembl protein sequences, Ensembl version 84 was used [33].

### Variation Data

The 1,000 Genomes Project phase 3 call set (02/05/2013 release) [2] was filtered to contain only single nucleotide variants using VCFtools, version 0.1.13 [34]. Variant calls were annotated using ANNOVAR, version 12/11/2014 [35], and GRCh37/hg19. For each protein a single isoform was used. Canonical protein isoforms, as identified by UniProt[32], were used where possible. If the canonical isoform was unknown, isoforms present in the Interactome3D data set were chosen first, otherwise the isoform was chosen numerically by its Ensembl protein identifier.

ANNOVAR was used to add functional annotation to variants. Corresponding UniProt and Ensembl protein sequences were aligned using NEEDLE, version 6.6.0 [36]. All protein sequence positions given in this study are in reference to the UniProt sequence, and variants that could not be mapped to UniProt sequence positions were excluded. ANNOVAR annotates each variant individually and does not consider multinucleotide variants. For each genome, we considered multiple variants in the same codon as a single change instead of two individual changes.

The gnomAD exome VCF dataset was downloaded on 26/04/2017, and the gnomAD allele frequencies of all variants from the 1,000 Genomes Project genomes were extracted [6,37].

### Protein Structure Data

All experimental protein structures were downloaded from the Protein Data Bank (PDB), and the Structure Integration with Function, Taxonomy and Sequence (SIFTS) database was used to map the residues in the protein structures to the residues in the protein sequences in the UniProt human proteome set [38,39]. Corresponding UniProt proteins and PDB structures were then aligned using MUSCLE [40]. Structures were selected for each protein, first prioritising resolution, and if two structures for overlapping sequence regions both had resolutions <3.5Å the structure with the largest sequence coverage was selected. After this step, all sequence positions covered by an experimental structure have been assigned to the highest quality structure available. For the remaining proteins without experimental structures, and for proteins with partial experimental structures where there are contiguous regions of ≥50 residues without a corresponding experimental structure, structural modelling was performed.

For structural modelling, template structures were identified from a 70% identity representative set of the PDB using hhblits, selecting only high-confidence templates with an hhblits probability ≥80 [38,41]. Models were selected to cover as many sequence positions as possible in a non-redundant manner, starting from the highest confidence model and selecting additional models that cover ≥50 sequence positions not already covered. Models were then generated for each sequence using the identified template and a protocol based on Phyre2 [42], with pulchra used to add and optimise the side chains [43]. Overall, combining multiple non-overlapping structures per protein and combining structural modelling with experimental structures greatly increased protein sequence coverage (Figure S1).

### Identification of Variant Combinations Within Proteins

Combinations of variants were identified within proteins for each of the 2,504 individual genomes, with each sample having between zero and two variant combinations per protein (one combination per allele). These variant combinations therefore consist of all of the variants that each sample has per copy of the protein, and they can be distributed anywhere within the 3-dimensional structure of the protein - termed Global Combinations (Figure 1.A). A second set of combinations was created for variants close in 3-dimensional space within structures - termed Proximal Combinations (Figure 1.A). All variants in Global Combinations were mapped to structures, and the distances between variants mapped to the same structures were calculated. Variants within 5Å of one another were grouped in to new combinations (Figure 1.A). For combinations containing >2 variants, 1 or more variants may be >5Å apart, providing each variant is within 5Å of another variant, and the maximum distance within the combination is <15Å (Figure 1.A). The vast majority of Proximal Combinations have a maximum distance between any two variants in the combination of 0-1Å (68.8% for non-synonymous variants). PyMOL (https://pymol.org/2/) was used to visualise variant combinations mapped to structures. Where described, variant combinations are given as a comma-separated list of variants, in ascending order by protein sequence position.

**Figure 1:**
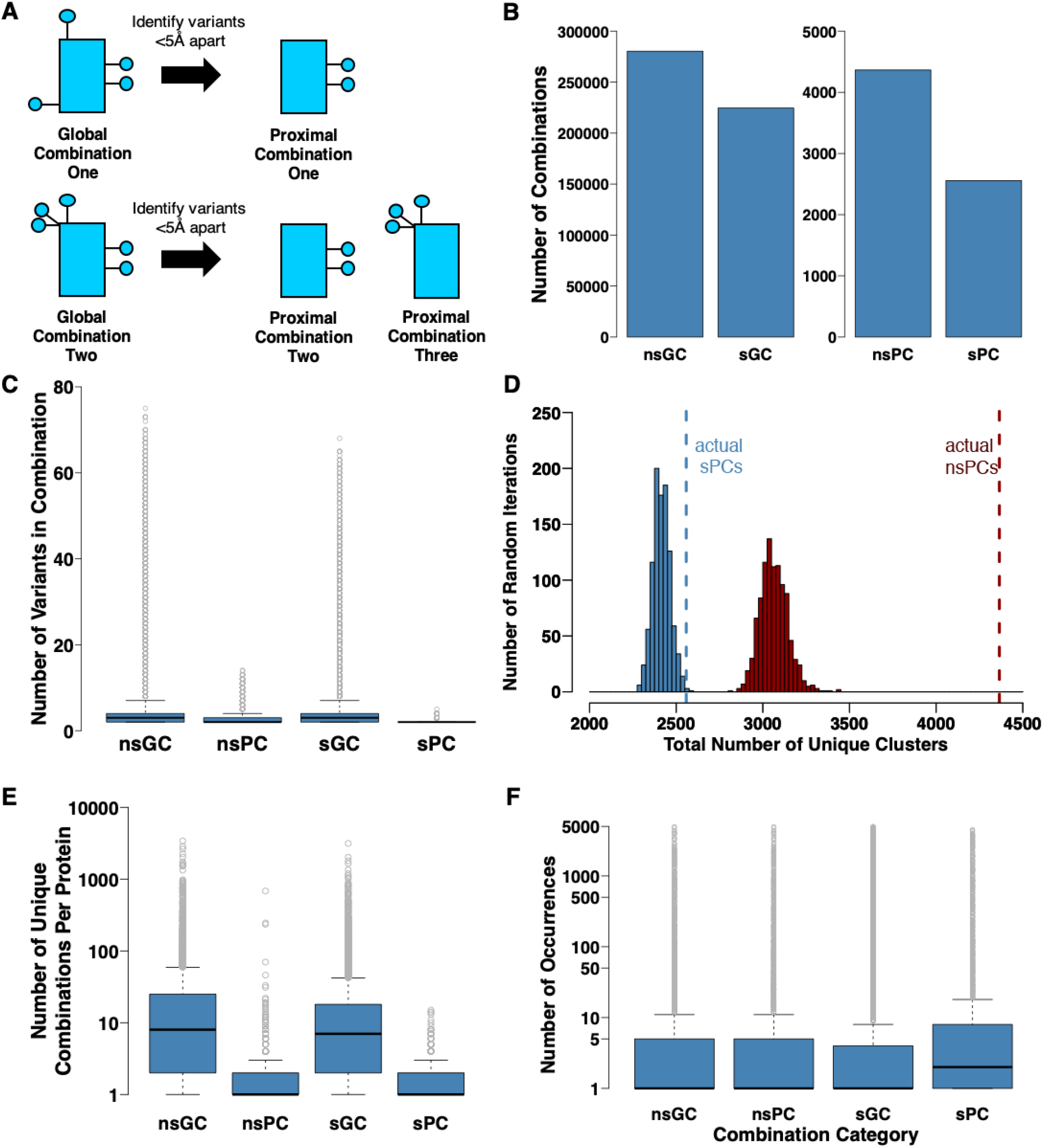
Global and Proximal Variant Combinations. The following abbreviations are used in the figure: nsGC – nonsynonymous Global combinations, sGC – synonymous Global Combinations, nsPC – nonsynonymous Proximal Combinations, sPC – synonymous Proximal Combinations **A)** Overview of Global and Proximal Combinations. **B)** Numbers of unique variant combinations **C)** Numbers of variants within combinations. **D)** Comparison of observed Proximal Combinations to random iterations. Bars represent the distributions of numbers of combinations for random iterations and the dashed lines represent the observed numbers of Proximal Combinations. Shown in Red for non-synonymous Proximal Combinations and blue for synonymous Proximal Combinations. **E)** Unique variant combinations per protein. **F)** Numbers of occurrences of each variant combination.

### Generation of Random Proximal Combinations

The positions of variants mapped to structures from Global Combinations were randomly assigned to different positions in the proteins, with the total number of variants in each protein kept the same, and Proximal Combinations were then redefined. For both non-synonymous and synonymous Global Combinations, 1,000 iterations were performed, creating 1,000 sets of Proximal Combinations for variants assigned to random sequence positions.

### Variant Property Compensations

Amino acid properties were assigned using the definitions of Innis et al., shown in Table S3 [44]. The sum changes in amino acid properties within variant combinations were then calculated to identify compensation of amino acid, charge, or functional group. Charge and functional group compensations were not counted if they were caused by a direct amino acid compensation, e.g. a direct compensation of an arginine residue is also a positive charge compensation, but would be counted only as a direct amino acid compensation. Foldx [45] was used to predict protein structure stability changes for variant combinations. The Foldx RepairPDB function was run for each protein structure with default parameters, and stability predictions were made using the BuildModel command on the repaired structures with default parameters.

### Interactome3D Data Processing

Protein-protein interaction data was taken from the Interactome3D database[46], Homo sapiens representative dataset version 2017_01[32]. Experimental structures and global templates were included, with domain-domain models excluded. The final set contained 9,642 distinct complexes (2,987 homomeric interactions, 6,655 heteromeric interactions, and 5,500 distinct proteins).

For each protein-protein complex, any residue in one partner within 5Å of a residue in the other partner of the complex was classed as being an interface residue[47]. Positions in structures were mapped to positions in UniProt sequences using MUSCLE [39].

### Identification of Interface Variant Combinations

Using the processed Interactome3D data, variants were classified into three different groups: interface, non-interface, and variants without interaction data. For each of the 2,504 genomes, interface variants were further classified in to one of four variant combination categories: Homomeric Combinations, Heteromeric Combinations, Uni-Partner Combinations, and Singletons. Where described, combinations of variants in interfaces use the following notation: <Protein A UniProt ID>:<Comma-Separated List of Variants in Protein A>_<Protein B UniProt ID>:<Comma-Separated List of Variants in Protein B>, e.g. PA:G21S,R43K_PB:P100H,H403P meaning that, for the interaction PA:PB, G21S and R43K variants both occur in the interface site of PA, and P100H and H403P both occur in the interface site of PB.

## RESULTS

### Global combinations of variants are common, while proximal combination are rare

Initially, combinations of both non-synonymous and synonymous variants in the 1000 genomes data were identified from the 531,560 non-synonymous and 345,334 synonymous variants present across the 2,504 genomes (see methods). For Global Combinations (multiple variants occurring in the same copy of a gene in an individual) a total of 280,329 non-synonymous Global combinations across 10,753 proteins and 224,541 synonymous Global Combinations across 11,131 different proteins were identified. The remaining variants occurred on their own without any further missense variants present in the same gene (342,031 nonsynonymous variants and 215,576 synonymous variants).

Global Combinations were filtered to identify Proximal Combinations, where variants occur close in 3-dimensional protein structure (see Methods) on the basis that such variants are more likely to be examples of evolutionary couplings. Variants that are distant in space may also coevolve via allosteric effects [48], but it is difficult to identify such variants. Proximal combinations were found to be much less common than Global Combinations, with 4,365 unique non-synonymous Proximal Combinations (across 1,562 proteins; Table S1) and 2,558 unique synonymous Proximal Combinations (across 1,678 proteins; Figure 1.B). Global Combinations of non-synonymous and synonymous variants on average contain more variants than Proximal Combinations (4.8 vs 3.4 variants for non-synonymous combinations, and 4.1 vs 2.1 for synonymous combinations). The numbers of variants in Global Combinations can be very high, with as many as 75 variants in a single combination, whereas the largest observed proximal combination contained 14 variants (Figure 1.C).

### Proximal Combinations Occur More Frequently than Expected by Chance

If Proximal Combinations are the result of coevolution between residues, we would expect to see more Proximal Combinations than if variants were randomly distributed throughout the protein. To test this, all variant positions were randomised on the protein structures 1,000 times and the number of random Proximal Combinations were calculated (see Methods). For non-synonymous variants, the number of Proximal Combinations observed in the 100 genome data is far greater than those obtained for random positioning of variants (42.4% more than the average from random positioning; 4,365 observed combinations, 3,064.61 mean random combinations and 3,429 maximum). This supports that there is a selective pressure for the variants in Proximal Combinations to be close in space and that they represent possible evolutionary couplings (Figure 1.D).

For synonymous variants, the number of Proximal Combinations falls within the distribution of the random positions (Figure 1.D), although the observed number is at the high-end of the distribution (2,558 observed combinations, 2,415.93 mean random combinations and 2,576 maximum). This may reflect that synonymous variants are unable to functionally compensate each other in the protein structure, but could be clustered in space due to sequence proximity, where any selection pressure is likely related to binding motifs in the DNA/RNA or selection of codons for translation speed related to protein folding [49-56].

The Proximal Combinations from the random iterations of non-synonymous variants also contained fewer variants on average compared to those from the sample data (mean 2.2 variants per combination vs 3.4). The maximum sizes of combinations from random iterations were also smaller, with an average value of 7.9, almost half the size of the maximum combination size from the variant data (14).

### Variant Combinations Per Protein

The unique variant combinations for each of the 20,791 proteins in the UniProt human proteome were considered (see Methods). Each protein has a distinct pattern of Global and Proximal Combinations, with some proteins having a large number of different combinations, and some having none or very few (Figure 1.E). The number of Combinations per protein is correlated with the length of the protein (*r*=0.54, Pearson’s product-moment correlation coefficient), but there is no correlation between Proximal Combinations per protein and protein length (r=-0.01), structural coverage (r=0.01), or numbers of residues within 5Å (r=0.01; Figure S2).

For non-synonymous variants, the average number of Global Combinations per protein was 26.07, and 20.17 for synonymous variants (Table 1), reducing to 2.79 and 1.52 for Proximal Combinations respectively (Figure 1.E). There are some proteins with a large number of Proximal Combinations, but these are rarer (Figure 1.E, Table 1), with the largest number observed in HLA class I histocompatibility antigen, B-81 alpha chain (684 unique non-synonymous proximal combinations; UniProt accession: Q31610). This protein functions in the immune system and many of the Proximal Combinations occur in the peptide-binding groove (Figure S3), a region of the protein recently reported to contain multiple pairs of residues that have coevolved in vertebrate evolution [60].

**Table 1:** Unique variant combinations observed per protein.

| Combination Category | Mean Number of Combinations | Median Number of Combinations | Maximum Number of Combinations |
| --- | --- | --- | --- |
| Non-Synonymous Global Combinations Per Protein | 26.07 | 8 | 3,402 |
| Non-Synonymous Proximal Combinations Per Protein | 2.79 | 1 | 684 |
| Synonymous Global Combinations Per Protein | 20.17 | 7 | 3,143 |
| Synonymous Proximal Combinations Per Protein | 1.52 | 1 | 15 |

Many of the observed variant combinations in proteins are rare, with 173,707 Global Combinations and 2,370 Proximal Combinations of non-synonymous variants only occurring in one heterozygous sample within the 2,504 genomes (Figure 1.F). In contrast some combinations are common, with 1,484 Global Combinations and 64 Proximal Combinations of non-synonymous variants with ≥1,000 occurrences (Figure 1.F). Some of these very common combinations reflect positions in the reference genome that do not represent the most common allele observed in human populations. An example of this is the combination ‘P47944:Y30C,W31R’ in protein Metallothionein-4, which occurs 4,398 times in the 1000 genomes data (Figure 2.I), while the allele counts in gnomAD [37] are 230,457 and 230,953 respectively.

gnomAD [37], a large aggregated resource of human sequencing data, was used to investigate the frequency of the variants occurring in combinations identified in the 1,000 genomes dataset. While gnomAD contains 123,136 exomes, which is many more individuals than the 2,504 in the 1,000 Genomes Project, as the aggregated it is not possible to identify variant combinations within individual genomes. However, this larger dataset was useful for looking at similarities in the overall frequencies of variants. For each Proximal Combination, the maximum allele count in gnomAD (i.e. most common variant in a combination) was compared to the mean allele count for the combination (Figure 2.A). The closer the allele count for the different variants in a combination, the smaller the difference between the maximum count and the mean count.

**Figure 2:**
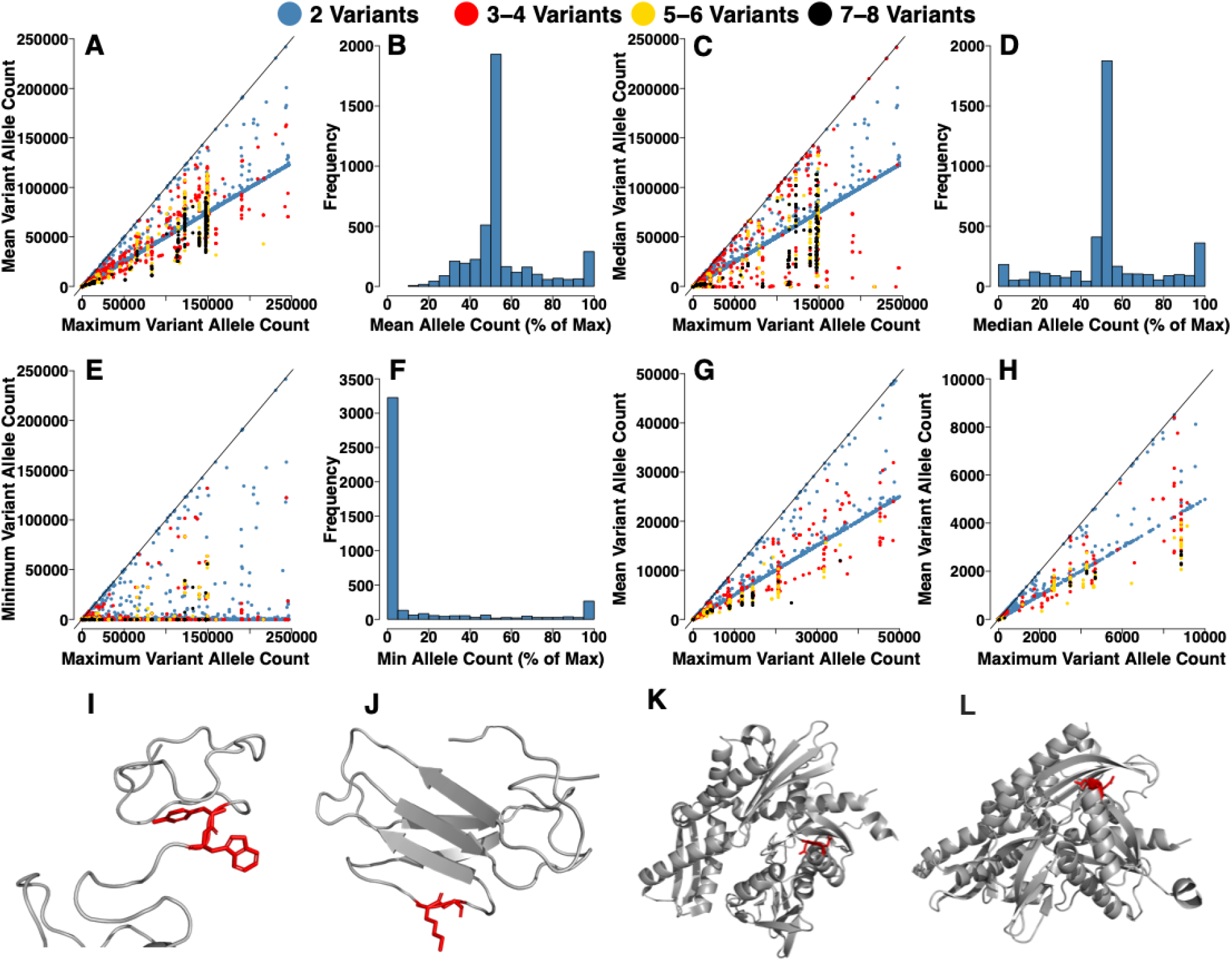
Allele counts of variants in gnomAD for the observed Proximal Combinations. In **A**,**C**,**E**,**G&H** each point corresponds to a variant combination, coloured by the number of variants in the combination, see legend. **A)** Maximum vs mean allele count within combinations. **B)** Distribution of the mean allele count as a percentage of the maximum allele count within combinations. **C)** Maximum vs median allele count per combination. **D)** Distribution of the median allele count as a percentage of the maximum allele count within combinations. **E)** Maximum vs minimum allele count per combination. **F)** Distribution of the minimum allele count as a percentage of the maximum allele count within combinations. **G)** Maximum vs mean allele count per combination, where maximum count <50,000. **H)** Maximum vs mean allele count per combination, where maximum count <10,000. **I)** Combination ‘P47944:Y30C,W31R’. **J)** Combination ‘A0A087WSZ9:K76M,G77R’. **K)** Combination ‘Q15109:W271R,C301S’. **L)** Combination ‘P17066:N153S,D154N’. Note - x-axes differ in their scales between the subplots.

A clear set of combinations have similar mean and maximum allele counts (points close to the x=y line in Figure 2.A) showing that there are variant combinations where the frequencies of all the variants are similar. There are 149 Proximal Combinations of non-synonymous variants where the mean allele count is identical to the maximum allele count, rising to, 352 and 476 where the mean count is >90% and >80% of the maximum count respectively. These are predominantly combinations containing two variants (blue points in Figure 2.A), although some larger combinations have similar mean and maximum allele counts. At lower maximum allele counts (below 50,000) there are combinations containing three or four variants (six combinations with mean count ≥90% of maximum count and 22 combinations ≥80%) where the mean is close to the maximum allele count (Figure 2.G&H). An off-diagonal line of points (gradient ∼0.5) is also observed, caused by pair combinations where one variant is common but the other variant is rare, resulting in a mean allele count approximately half of the maximum allele count (Figure 2.A&B).

Comparison of the maximum and median allele counts (Figure 2.C&D) identified a set of combinations (mainly containing three or four variants; red points in Figure 2.A), where the median allele count is much closer to the maximum allele count than the mean allele count. These represent combinations where a subset of the variants have similar frequencies but the remaining variants occur rarely (e.g. a combination containing three variants where two have similar frequencies and a third variant that occurs in a single individual). This could represent scenarios where these two positions in a combination have coevolved but a third variant is occasionally also present. There are 157 Proximal Combinations of non-synonymous variants where the median allele count is identical to the maximum allele count, rising to 267, 452 and 638 Proximal Combinations where the median count is >99%, >90% and >80% of the maximum count respectively. These numbers are greater than when compared to the mean allele count (see above) suggesting that some subsets of variants occur frequently together, while the remaining variants in the combination do not.

Further, this is supported by the minimum allele counts, for combinations with greater than two variants the minimum allele count is almost always much lower than the maximum allele count (Figure 2.E&F). This indicates that for most of these combinations containing >2 variants it is rare for all variants to have similar frequencies. There are 149 Proximal Combinations of non-synonymous variants where the minimum allele count is identical to the maximum allele count, rising to 207 where the minimum count is >99% of the maximum count. This further demonstrates Proximal Combinations where variants are nearly always observed together.

For 153 of the observed Proximal Combinations the allele counts of the constituent variants are identical in gnomAD (with 30 of these having allele counts ≥25), and it is more likely for these variants to have a strong coevolutionary relationship. For example, in Heat shock 70 kDa protein 6 (UniProt accession: P17066), the combination ‘P17066:N153S,D154N’ (a potential direct amino acid compensation of asparagine) occurs 63 times (Figure 2.K), and both variants have allele counts of 561 in gnomAD. In other proximal combinations the allele frequencies are very close, such as for the combination ‘A0A087WSZ9:K76M,G77R’ in the protein T cell receptor alpha variable 30, which occurs 283 times in the 1000 genomes data set (Figure 2.J), and the variants have allele counts in gnomAD of 7,479 and 7,477 respectively. This example represents variants where we see charge compensation (loss of lysine and gain of arginine – see below).

Despite the gnomAD data being aggregated, in combination with the 1,000 Genomes Project data it provides additional support for variants with a likely coevolutionary relationship.

### Proteins have a Predominant Proximal Combination

Proteins can have multiple different variant combinations present in individuals, sometimes with overlapping variants. However, for Proximal Combinations, most proteins have a predominant variant combination, that accounts for the majority of variant combinations observed (Figure 3.ii; i.e. the most common Proximal Combination accounts for more than 50% of the variants observed), even for proteins where there are a large number of genomes with variant combinations in the protein (points on the right of the x-axis in Figure 3.ii). Of the 1,562 proteins with non-synonymous Proximal Combinations, 1,098 (70.3%) have a combination that accounts for ≥90% of all combinations, however 544 of these only occur once in the 2,504 individuals. In contrast, only 26.9% of proteins with non-synonymous Global Combinations have a combination that accounts for ≥90% of all combinations (Figure 3.i).

**Figure 3.**
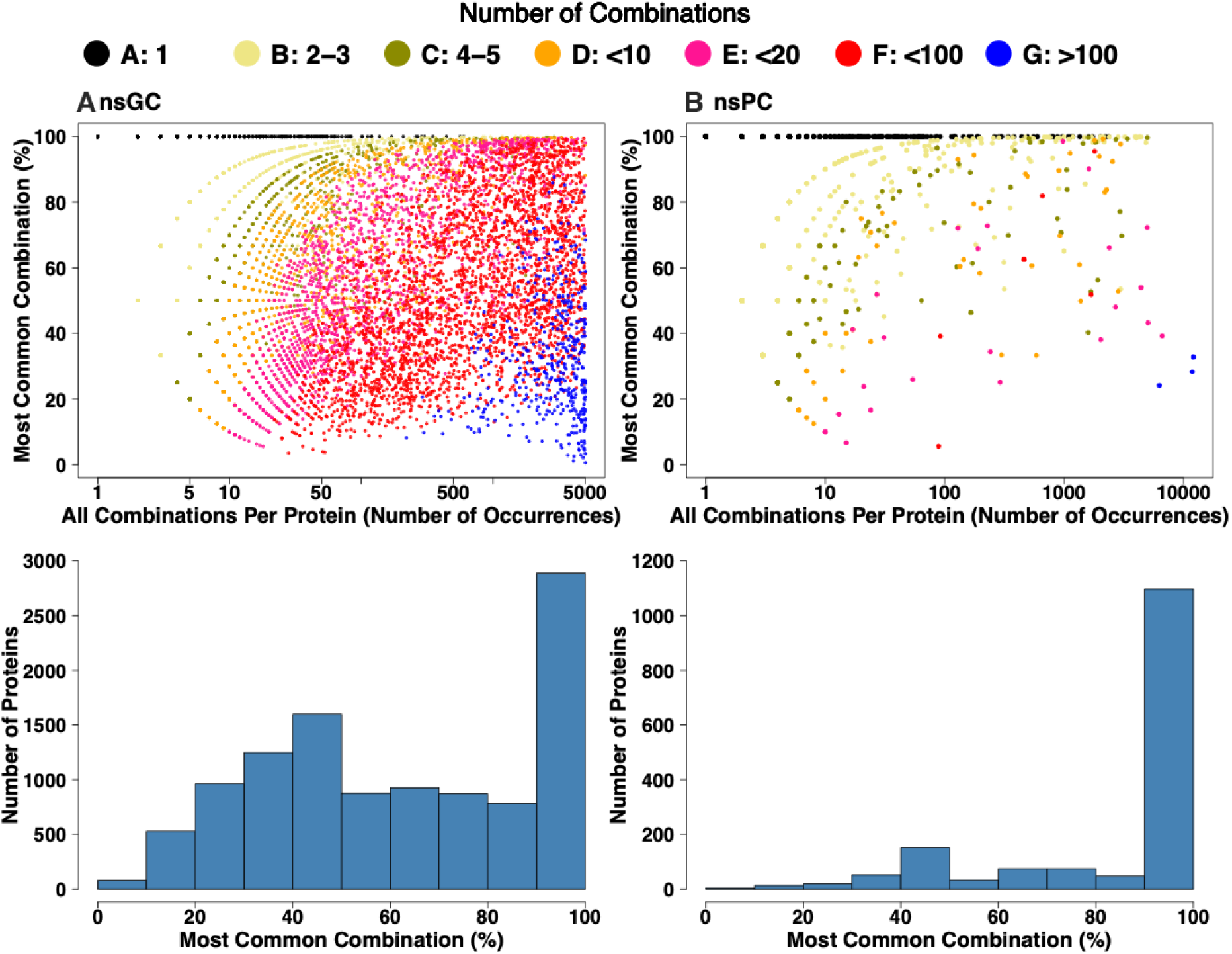
Comparison of most common variant combinations in gnomAD. Occurrences of the most common variant combinations vs all variant combinations in individual proteins. Each point is a protein, the colour is determined by the number of combinations observed in the protein, see legend. The x-axis position is determined by the number of occurrences of any combination within the protein, and the y-axis position is determined by the percentage of total occurrences accounted for by the most common variant combination. If the most common variant combination accounts for all of the variant combinations in the protein (black points) the point will lie at the very top of the y-axis (100% of combination occurrences are the most common combination). The bottom sections show the distributions of proportions of variant combination occurrences from the most common variant combination per protein. **A)** Non-Synonymous Global Combinations. **B)** Non-Synonymous Proximal Combinations. Note - x-axes differ in their scales between the subplots.

While some combinations are spread relatively evenly across the super populations present in the 1000 genome dataset (Figure S4 - light blue boxes across the super populations). Others are concentrated in a subset of super populations (Figure S4 - dark blue boxes in specific super populations). For example, the Proximal Combination ‘G3V1Y8:C95R,L96V’ is most common in the African super population (AFR; 43.6%), whereas ‘Q8N423:H300Y,C306W’ is most common in the European super population (36.4%; Figure S4).

Larger differences in the population distributions of Proximal Combinations are seen for less frequent combinations (as common combinations are present in many individuals and therefore are likely to be present in multiple populations). We considered the 50 most common Proximal Combinations where the total number of combination occurrences is <500, many of these combinations are most common in the AFR super population (Figure S4), such as ‘A0A0B4J1V5:A91T,K93E’, and ‘Q8WXQ8:L336S,S378G’ - a potential variant compensation of serine. The combination ‘Q9H339:T78K,R88G’ occurs 486 times in total, with 82.7% of these in the AFR super population, and is another example of a potential variant effect compensation (compensation of a positive charge, with loss of arginine and gain of lysine). There are also examples of combinations that are more common in one or more of the other four super populations, such as A6NGD5:G108S,V109M’, for which 57.6% of occurrences are from the South Asian (SAS) super population (Figure S4).

Variant combinations that are evenly distributed between the super populations are likely to have first occurred longer ago in human evolution. Population-specific variant combinations are likely to have occurred more recently, and there are individual combinations that are more common within each of the super populations. For each super population there are thousands of combinations where >90% of the total occurrences are from that super population, but the AFR super population clearly has many more of these combinations than the other four super populations (Figure S5.A).

### Variant combinations frequently show compensation of properties

Coevolution occurs when a subsequent variant arises that compensates for a variant that is already present. To further identify possible coevolution (or evolutionary couplings) of the Proximal Combinations of variants, we considered if the variants could compensate for each other. Direct amino acid compensations are the simplest and most complete type of compensation event. For example, in a pair of variants one position changes from alanine to serine and the second position changes from serine to alanine, such that the same amino acids are present but in different positions in the structure. This is the case for the Proximal Combination ‘A6NLU0:S276A,A277S’ in the protein Ret finger protein-like 4A. This combination occurs 1,186 times in total, predominantly in the African and South Asian super populations and these variants are rarely observed on their own (S276A twice and A277S once). For the 4,365 Non-Synonymous Proximal Combinations, 571 unique direct amino acid compensations were observed (Figure 4.A). Some individual amino acids are compensated for more often than others, with arginine compensations occurring most frequently (85, 14.8% of total) (Figure 4.B), followed by serine (64 - 11.2%). The level of direct amino acid compensation is correlated with the number of codons per amino acid (*r*=0.77; Figure S2.i).

**Figure 4:**
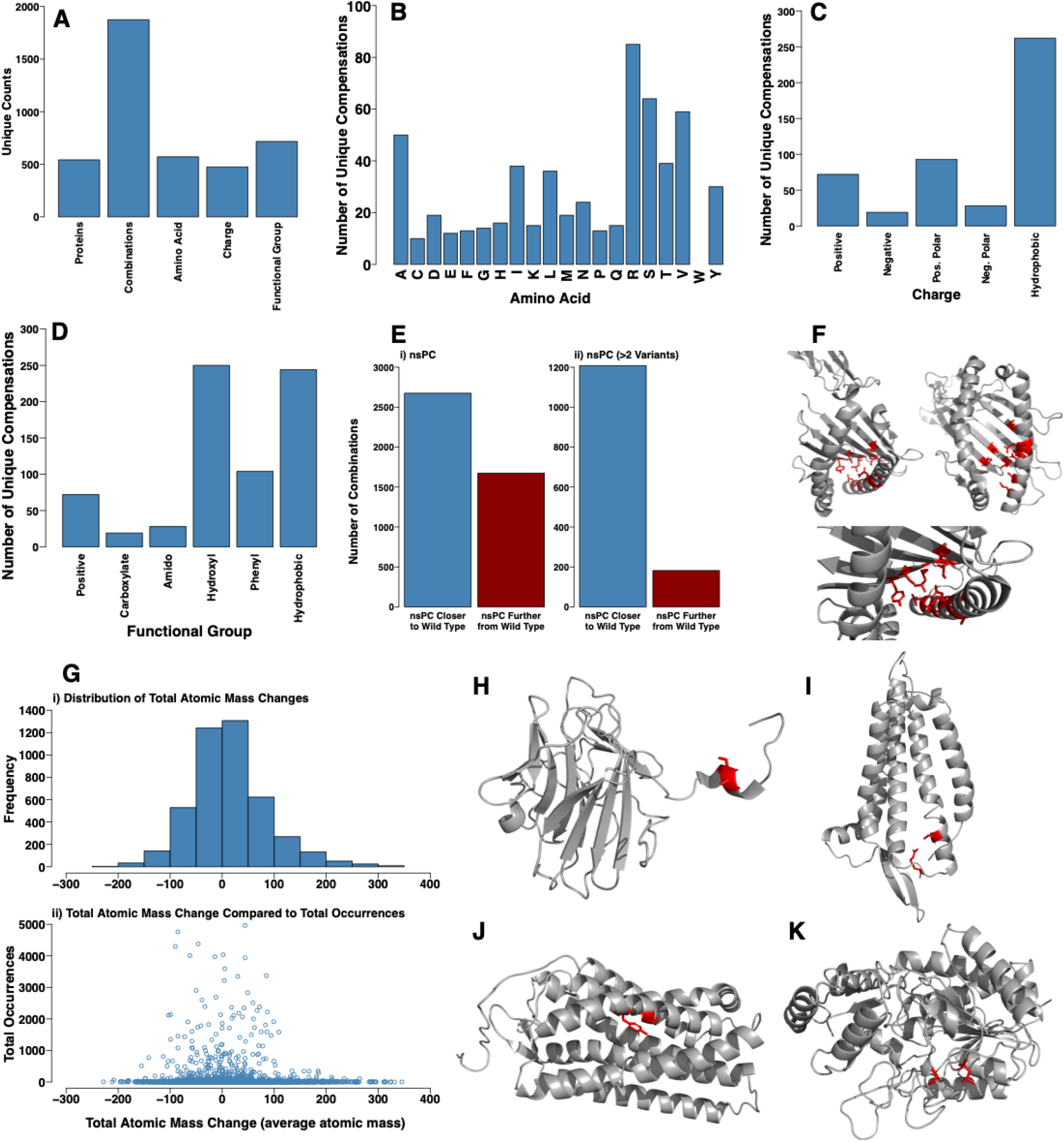
Compensatory effects within Proximal Combinations. **A)**. Unique occurrences of functional group compensations **B)** Unique occurrences of direct amino acid compensations. **C)** Unique occurrences of charge compensations. **D)** Numbers of property compensations. **E)** Foldx stability change prediction distributions for combinations containing two variants. i) Single variants within the combinations. ii) Both variants together. **F)** Foldx stability change prediction distributions for combinations containing >2 variants. i) Single variants within combinations. ii) All sub-combinations for the observed variant combinations. iii) Full variant combinations. **G)** Absolute stability change prediction values for full combinations vs sub-combinations with the largest predicted change in stability. i) All combinations. ii) Combinations containing >2 variants. **H)** The combination ‘Q31610:S35A,V36M,S48A,A93T,Q94N,A95T,D98Y,S101N, L105A,L119W, S121T,Y140F’. **I)** Total mass changes for combinations. i) Distribution of mass changes. ii) Total mass change vs occurrences of each combination. **J)** Potential direct amino acid compensation in ‘A6NLU0:S276A,A277S’. **K)** Potential positive charge compensation in ‘Q9H339:T78K,R88G’.

Potential compensation of charge (474) and amino acid side chain functional groups (717) were also identified (Figure 4.A; see Methods). Functional group compensations were the most common type of compensation observed, followed by direct amino acid compensations, while charge compensations were the least common (Figure 4.A). When considering compensation of charge, hydrophobic compensations were the most common (262 - 55.27% of all charge compensations; Figure 4.C), likely reflecting the larger number of hydrophobic sidechains.

Charge compensation is observed in the previously discussed combination ‘Q9H339:T78K,R88G’ (82.7% of the 486 occurrences within the AFR super population), where a positive charge is lost in the R88G variant, but a positive charge is gained in the T78K variant, which maintains a positively charged residue in this region of the protein structure (Figure 4.I). Q9H339 corresponds to the protein Olfactory receptor 51B5, which is involved in the sensory perception of smell. Both T78K and R88G occur in an extracellular domain of the protein, therefore these residues may be involved with ligand binding necessary for olfactory perception, and loss of the positively charged arginine residue without compensation by the gain of the positively charged lysine residue could result in altered ligand binding. Across the data set, there is only one occurrence of T78K without R88G, and only 42 occurrences of R88G without T78K.

For functional group compensations (positive, carboxylate, amido, hydroxyl, phenyl, and hydrophobic), hydroxyl (250 unique; 34.86% of total) and hydrophobic (244 unique; 34.03% of total) are most frequently observed (Figure 4.D). In the protein Olfactory receptor 7G1 (UniProt accession: Q8NGA0), the combination ‘Q8NGA0:S249F,Y252C’ occurs 139 times (Figure 4.J), and is an example of a potential functional group compensation – with one phenyl group lost with the variant Y252C potentially compensated for by the gain of another phenyl group with S249F. Interestingly, in this example two hydroxyl groups are lost within close spatial proximity without replacement.

Variant combinations can involve multiple compensation types. The previously discussed combination ‘Q8WXQ8:L336S,S378G’ in the protein Carboxypeptidase A5 is an example of a potential direct amino acid compensation and a charge/functional group compensation (Figure 4.K). In this example, serine is lost and replaced and a small hydrophobic sidechain is also retained. This combination occurs 273 times, and predominantly occurs in the AFR super population (88.64% in AFR).

Overall, a potential functional compensation involving direct compensation of one or more amino acids, compensation of one or more charge types, or compensation of one or more functional groups was identified for 1,875 Proximal Combinations across 542 proteins. This corresponds to potential compensatory effects of variants in 42.96% of the 4,365 total Proximal Combinations identified and highlights that many of the Proximal Combinations are likely to represent evolutionary couplings (Figure 4.A).

### Full Variant Combinations Are More Stable Than Sub-Combinations

Foldx [45] was used to analyse the effect of variant combinations on protein stability as compensation may occur to maintain protein stability (see methods). This considered if the full combination of variants was predicted to have a smaller effect on protein stability than a sub-combination. For example, in Q31610 (Figure S3) the combination ‘Q31610:S35A,V36M, S48A,A93T,Q94N,A95T,D98Y,S101N,L105A,L119W,S121T,Y140F’ was predicted to destabilise the protein (+4.53 kcal/mol; Figure 4.F), while sub-combinations were predicted to be far more destabilising e.g. (‘Q31610:S11A,A69T, Q70N,A71T,D74Y, S77N,L95W’ +19.21 kcal/mol) and thus the full combination may compensate for this.

Overall, the majority (2,684; 61.5%) of full combinations were predicted to be closer to the wild type stability of the protein than one or more sub-combination of variants, rising to 86.9% for Proximal Combinations containing more than two variants (Figure 4.E). Therefore, for the majority of Proximal Combinations there are sub-combinations of variants that are predicted to cause a larger change in structural stability (stabilising or destabilising), indicating possible selection of combinations of variants that preserve the wild type protein stability, which is more pronounced for combinations with >2 variants, (Figure 4.E.ii), providing further evidence for potential evolutionary couplings.

Variant compensation can also account for changes in the size and shape of amino acid sidechains. Amino acid mass was used as a proxy to consider these properties (it captures the size of the sidechain but not the structure). The majority of combinations have an atomic mass change between +/-50 (58.4%), rising to 84.8% within +/-100 (Figure 4.G), suggesting that many combinations compensate for mass. Some combinations show large reductions (greatest reduction -228.3) or increases (+345.4 greatest increase) in atomic mass, however, such combinations occur rarely in individuals (Figure 4.G), while combinations that occur frequently in the 1,000 genomes data set largely have a mass change within +/-50 (Figure 4.G).

### Compensating Combinations Also Occur in Protein-Protein Interfaces

Protein-Protein interactions are essential for cellular function, a key component of cellular complexity, and their interface sites are enriched in disease associated variants [55-58]. We therefore expanded our analysis to identify variant combinations occurring within interface sites. Using interaction data for 9,642 protein-protein interactions with residue-level structural characterisation from Interactome3D [46] we identified 6,824 unique non-synonymous variants present in interface sites, 592 of which form at least one variant combination (Table S2). Spread across 233 proteins and 272 complexes there were 147 Homomeric, 364 Heteromeric, and 735 Uni-Partner combinations (where there were multiple variants in the interface of a single protein within a heteromeric complex; Figure 5.A), reflecting that the majority of complexes in Interactome3D are heteromeric (Figure 5.B; see methods).

**Figure 5:**
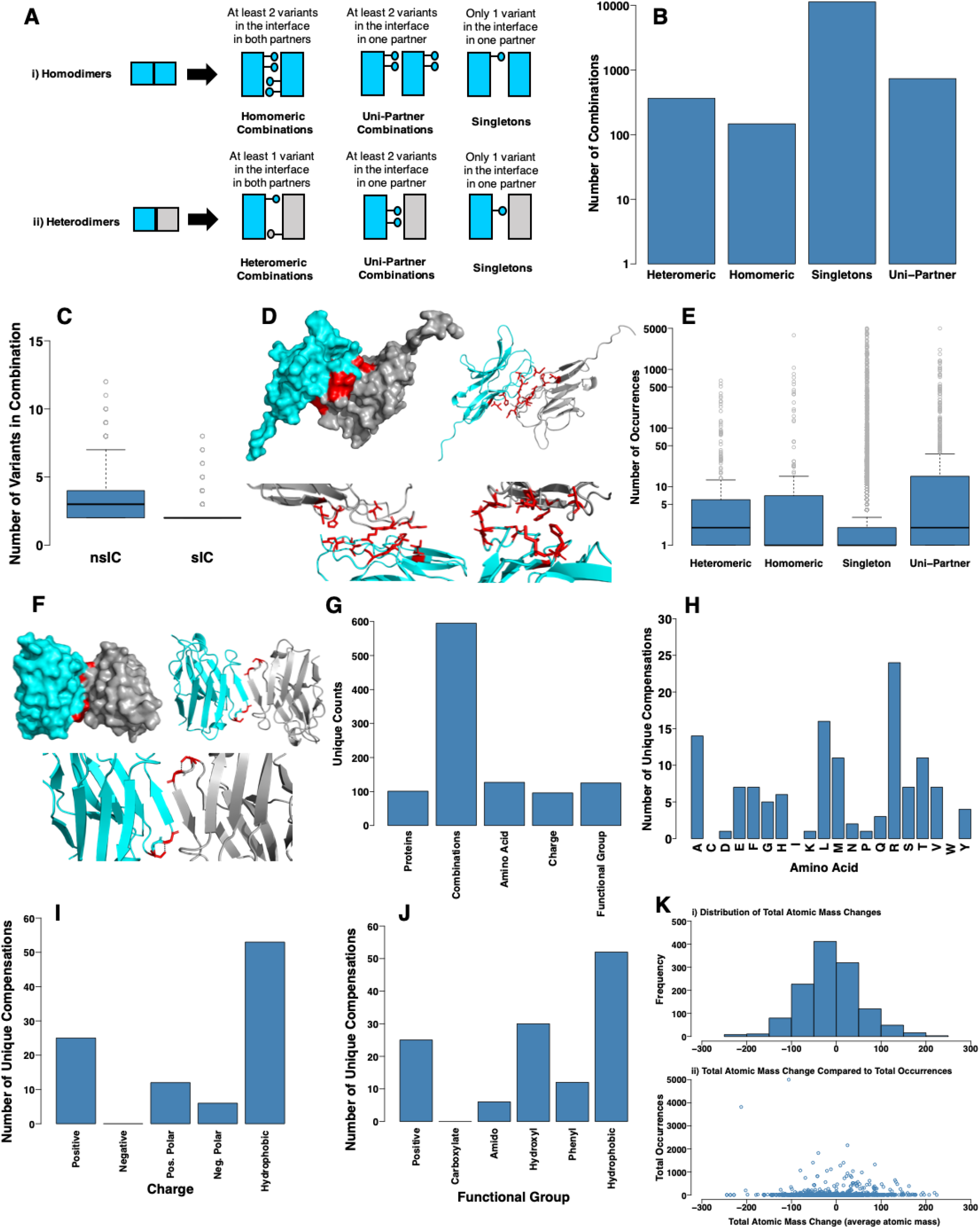
Interface variant combinations. **A)** Overview of Interface Variant Combinations. **B)** Numbers of interface variant combinations observed. **C)** Numbers of variants within combinations. **D)** Combination ‘P78324:T52I,R54H,A57V,D95E,L96S,N100K_P78324:T52I,R54H,A57V,D95E,L96S, N100K’. Chain A - grey, B - cyan, and positions of variants - red. **E)** Occurrences of each combination. **F)** Combination ‘P01764:S73G,G75S’. Chain A - grey, chain B - cyan, positions of variants - red, and the hydrogen bond between the hydroxyl group of S73 and the backbone of G75 - dashed black line. **G)** Numbers of potential property compensations within combinations. **H)** Occurrences of potential direct amino acid compensations. **I)** Occurrences of potential charge compensations. **J)** Occurrences of potential functional group compensations. **K)** Total mass changes for combinations. i) Distribution of mass changes. ii) Total mass change vs occurrences of each combination.

As with Proximal Combinations, where individual proteins had distinct patterns, different protein complexes were observed to have different patterns of interface variant combinations. The majority of the interface variant combinations contain only a few variants, with 525 combinations containing only two variants, but some are much larger, containing as many as 12 variants (Figure 5.C). For example, there are 209 occurrences in the homodimer of Tyrosine-protein phosphatase non-receptor type substrate 1 protein (SIRP-α; P78324) of an interface variant combination of the same six variants in both copies of the protein (Figure 5.D). SIRP-α is a cognate receptor for CD47, and the interaction of the two is involved in regulation of Interleukin-12 levels, possibly as a homeostatic mechanism to prevent escalation of the inflammatory immune response [61]. Multiple potentially compensatory effects are present in this set of variants. This includes R54H and N100K which results in a large positively charged sidechain being retained. While T52I and L96S retain a hydroxyl group and a hydrophobic sidechain. Finally, the variants D95E and A57V are both conservative changes, one maintaining a negative charge (D95E) and the other a small hydrophobic amino acid (A57V). Therefore, despite the occurrence of six different variants, the overall properties of the positions in the interface are largely maintained.

Overall, many of the interface combinations observed are rare (Figure 5.E) and for most complexes there is a predominant variant combination accounting for the majority of the total combinations in the interface (Figure S6), which for the majority of combinations accounts for >90% of all combinations in the complex. This holds across the three combination types; 64% of Heteromeric, 60% of Homomeric, and 73% of Uni-Partner Combinations.

The protein Immunoglobulin heavy variable 3-23 (UniProt accession: P01764) plays a role in antigen recognition, and forms a homomeric complex. In this complex the variant combination ‘P01764:S73G,G75S’ occurs 154 times (Figure 5.F). This is a clear example where the combination of variants could be compensatory and represent an evolutionary coupling. In both chains of the complex, serine is lost at position 73 and a glycine is gained, and then at position 75 a glycine is lost and a serine is gained. The wild type and variant type amino acids are balanced, with no net change in the number of serine or glycine residues. Neither of these variants were observed on their own in any of the 2,504 genomes, they only occur together. One possible advantage of the two variants occurring together, and neither alone, is maintaining the hydroxyl group of serine. In the wild type structure S73 acts as a hydrogen bond donor, forming a hydrogen bond with the backbone of G75 (Figure 5.F), which is likely stabilising the tight turn in the structure.

As with combinations of variants within individual proteins (Figures S4&S5.A), some of the interface variant combinations that are rare across the dataset are common within specific populations (Figure S5.B). Many combinations occur almost exclusively in the African super population for all three types of interface variant combination, with 267 combinations where >90% of the occurrences are in the African super population (72 for AMR, 66 for EUR, 108 for EAS, 96 for SAS). Many of these combinations are rare, and when filtering for combinations with ≥25 total occurrences this number falls to 22 for the AFR super population (0 for AMR, 0 for EUR, 3 for EAS, 0 for SAS). This likely reflects that the human reference genome is least representative of the AFR super population [62]. The previously discussed combination ‘P01764:S73G,G75S’ in P01764:P01764 (Figure 5.F), occurs in 149 individuals, 38.26% from AFR, 10.74% from AMR, 46.98% from EAS, 4.03% from SAS, and 0.00% from EUR.

In total, 595 combinations with compensations were observed across 101 unique interfaces (there may be multiple combinations per interface), representing 47.8% of interface combinations. These were comprised of 127 amino acid compensations, 96 charge compensations and 125 functional group compensations (Figure 5.G). Similar patterns of compensations were observed (as for those in individual protein structures – Figure 4.A) with arginine being the most commonly compensated amino acid (19% of direct amino acid compensations; Figure 5.H), and this shows some correlation with the number of codons per amino acid (*r*=0.62; Figure S2), but less so with the amino acid composition of interfaces (*r*=0.41; Figure S2). Compensation of hydrophobic sidechains were the most common charge and functional group compensations (Figure 5.I&J). For charge compensations, this is followed by compensation of positive charge, while hydroxyl group compensations were the second most common type of functional group compensation (24%). Total mass changes of interface variant combinations are on average fairly conservative (mean -11.0, median -12.1, min -244.4, max +224.176, 86.6% between -100 and +100 total mass change; Figure 5.K), similar to proximal combinations within structures (Figure 4.G).

### Variant Combinations Present in Individual Genomes

The combinations present in each individual genome were considered to identify the number of compensations present in a typical individual. Individuals have an average of 2,430 global combinations (range 2,115 - 2,882), 122 proximal combinations (range 69-177), and 25 interface variant combinations (range 4-64; Figure 6). Given that these genomes have between 10,000-12,000 non-synonymous variants and the human genome has approximately 20,000 genes, this puts the number of global combinations into context. Although genomes have a large number of proteins with multiple variants in them (Global Combinations) they have relatively few proteins with multiple variants close in space (Proximal Combinations). For interface variant combination categories, individuals have averages of 4.4 Homomeric Combinations (range 0-17), 2.8 Heteromeric Combinations (range 0-14), and 18.0 Uni-Partner Combinations (range 3-43; Figure 6.C).

**Figure 6:**
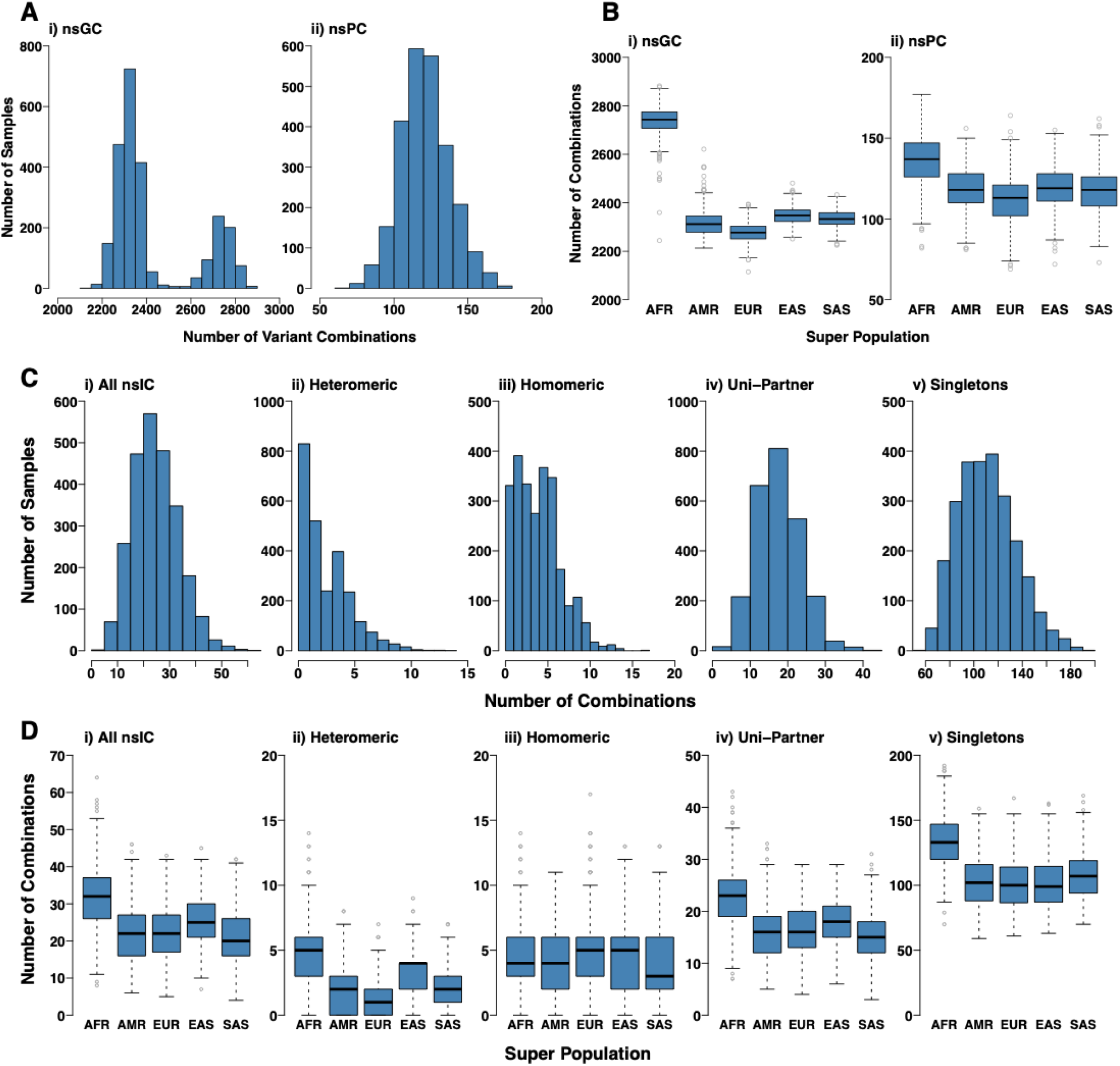
Variant combinations per genome. **A)** Variant combinations within structures. **B)** Variant combinations within structures per super population. **C)** Interface variant combinations. **D)** Interface variant combinations per super population. AFR – African super population, AMR – American super population, EAS – East Asian super population, EUR – European super population, SAS – South Asian super population.

The number of non-synonymous Global Combinations per individual exhibit a bimodal distribution (Figure 6.A.i), which is not observed for the Proximal Combinations (Figure 6.A.ii). This is caused by population-specific differences between the genomes, with those from the African (AFR) super population having more non-synonymous Global Combinations compared to the other four super populations (Figure 6.B.i), which results in the bimodal distribution of Global Combinations per sample observed (Figure 6.A.i). AFR genomes have on average the most Proximal combinations, but the difference between the super populations is much smaller than for the Global Combinations (Figure 6.B.ii). Genomes from the AFR super population also have on average more interface variant combinations than genomes from the other four super populations (Figure 6.D). This trend is observed for Heteromeric and Uni-Partner interface variant combinations, but not for Homomeric combinations (where genomes have similar numbers between the five super populations; Figure 6.D).

## DISCUSSION

In this study we sought to identify, examples of coevolution within human genomes (evolutionary couplings). To do this we quantified variant combinations in individual human genomes at three different levels: variant combinations across protein sequences (280,329 non-synonymous), variant combinations in close proximity in 3-dimensional protein structure (4,365 non-synonymous), and variant combinations within protein-protein interface sites (1,246 non-synonymous). These combinations show considerable variability in their distributions between populations, with some combinations distributed evenly across populations and others heavily skewed towards individual populations. At the protein level most proteins have one predominant combination of variants, especially for variant combinations in close spatial proximity. Others have begun to consider variant combinations within the 1,000 genomes data, for example, Spooner et al., [63] recently developed a resource to consider haplotypes for proteins that can be used to inform drug design. Here our focus is on performing the first genomic survey of coevolution within the human genome.

Our analysis demonstrated that Proximal Combinations occur much more frequently than expected by chance (Figure 1.D), indicating that such variants are not observed by coincidence. However, proximity in space does not infer that variants are coevolving. Our analysis shows that for 72.1% (3,145 of the 4,365) of Proximal Combinations there are compensations that result in direct change of amino acid, conserve charge or functional groups, or are predicted to preserve structural stability. This suggests that the majority of Proximal Combinations are likely to represent examples of evolutionary couplings. For 47.8% of interface combinations (595 of 1,246) potential compensations were also observed. This value is lower as it does not consider predicted stability changes but still shows that a significant proportion of interface combinations represent examples of evolutionary couplings.

The comparison with gnomAD [37] further supports our findings by identifying variants that occur with similar frequencies in a much larger dataset (123,136 exomes), unfortunately as the gnomAD data is aggregated it was not possible to use gnomAD for the main analysis that we have performed. However, we did identify a set of Proximal Combinations where the variants in gnomAD have identical or very similar allele counts.

The variant combinations that we have identified in this study are likely a fraction of the true number that co-occur within human genomes. Firstly, we have only considered protein coding changes (and only single amino acid variants), and there are undoubtedly combinations involving non-coding variants with compensatory effects. Secondly, for simplicity we only considered the canonical isoform of each protein. There will be variant combinations that only occur within non-canonical isoforms, although recent work has suggested that for most proteins there is only one functionally relevant isoform at the protein level [64]. Finally, Proximal Combinations can only be identified for proteins with an experimental or modelled structure. For the 20,791 proteins in the human proteome set used in this study, 4,138 have no structural data available, with the remaining proteins having an average coverage of 59.5% of their sequence (see Methods). Similarly, a structure for each protein-protein complex is required in order to determine interface variant combinations. The release of Interactome3D [46] used in this study contains 9,642 unique protein-protein complexes (see Methods), a small fraction of the >1,000,000 [65] protein-protein interactions predicted to constitute the human interactome.

Nevertheless, this study provides an important first, to our knowledge, analysis of coevolution that has occurred within the human genome. Our findings highlight the necessity for programs that predict the effect of single nucleotide variants [66-73] to consider the context in which these variants occur. Individually programs may predict such variants to be deleterious, however given that we observe many compensations for possible functional effects, such predictions would be incorrect. Therefore, it is essential that the next generation of methods consider variant effect prediction in a much broader sense, considering other variants in the genome, rather than each one individually. To date very few methods have made use of evolutionary couplings to predict the effect of nsSNVs [74]. Combinations of variants in individual human genomes will determine resilience or susceptibility to a host of selection pressures, from inherited disease, to drug response, to pathogenic infection. The interpretation of gestalt effects of variant combinations will be a key challenge in the future of precision medicine.

## Supporting information

All supplementary Figures and Tables

## DATA AVAILABILITY

All relevant data are available in the text and supplementary material.

## SUPPLEMENTARY DATA

Supplementary Data are available at NAR online.

## ACKNOWLEDGEMENT

We thank Martin Michaelis for helpful comments on the manuscript.

