## Supplementary material for "Individual human genomes frequently contain variants that have evolutionary couplings": All supplementary Figures and Tables

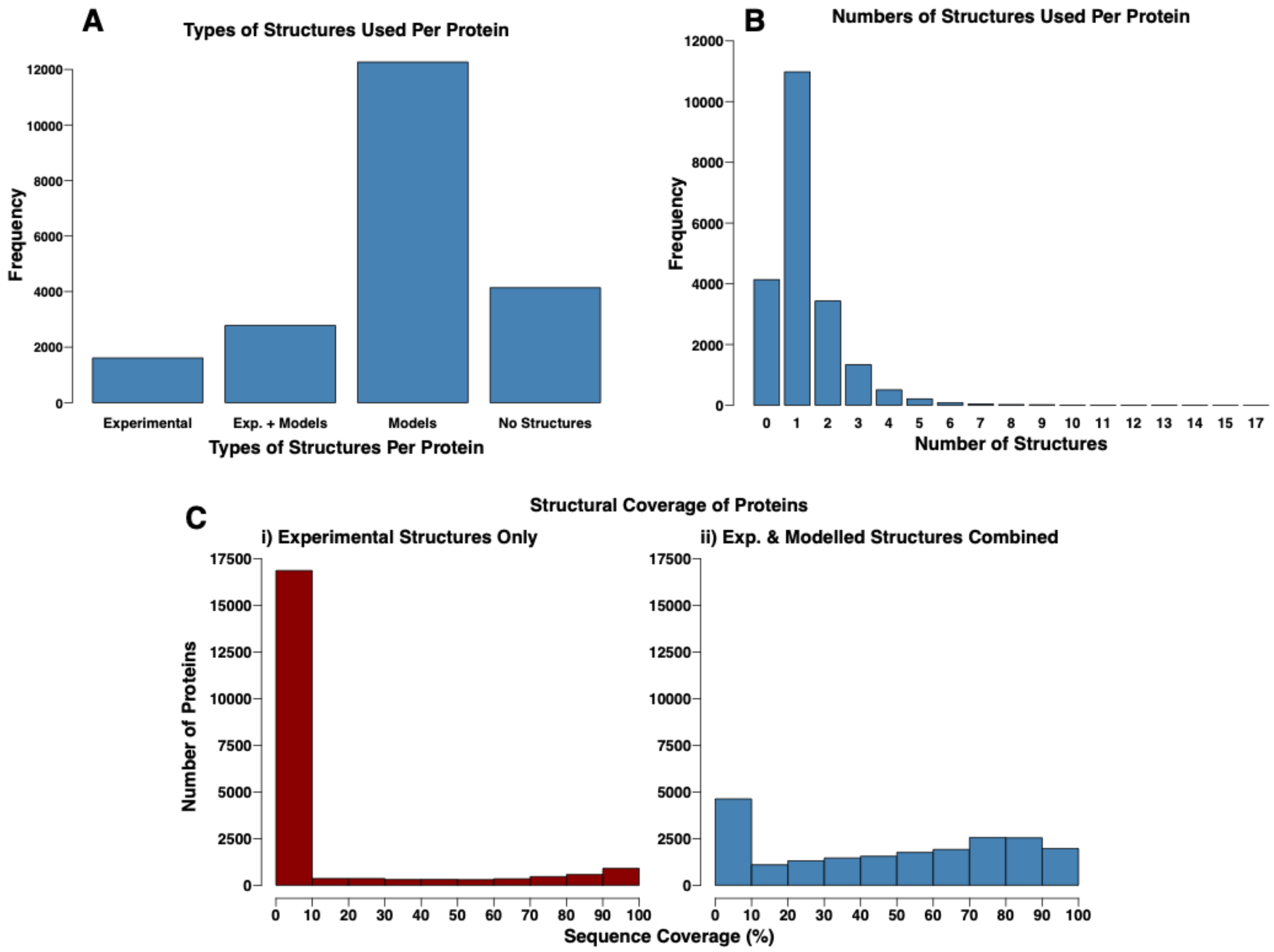


**Figure S1.** Protein structures used in the analysis. **A)** Types of structures used per protein. Experimental – proteins with corresponding experimental structures, with no sequence gaps ≥50 residues. Experimental + Models - proteins with corresponding experimental structures, but with sequence gaps ≥50 residues for which structural modelling was performed. Models – proteins for which there were no available experimental structures and structural modelling was performed. No structures – proteins without experimental models and for which no high-quality structural modelling template was identified. **B)** Numbers of non-overlapping structures used per protein. **C)** Percentage protein sequence coverage per protein. **i)** Using only experimental structures. **ii)** Using experimental and modelled structures.

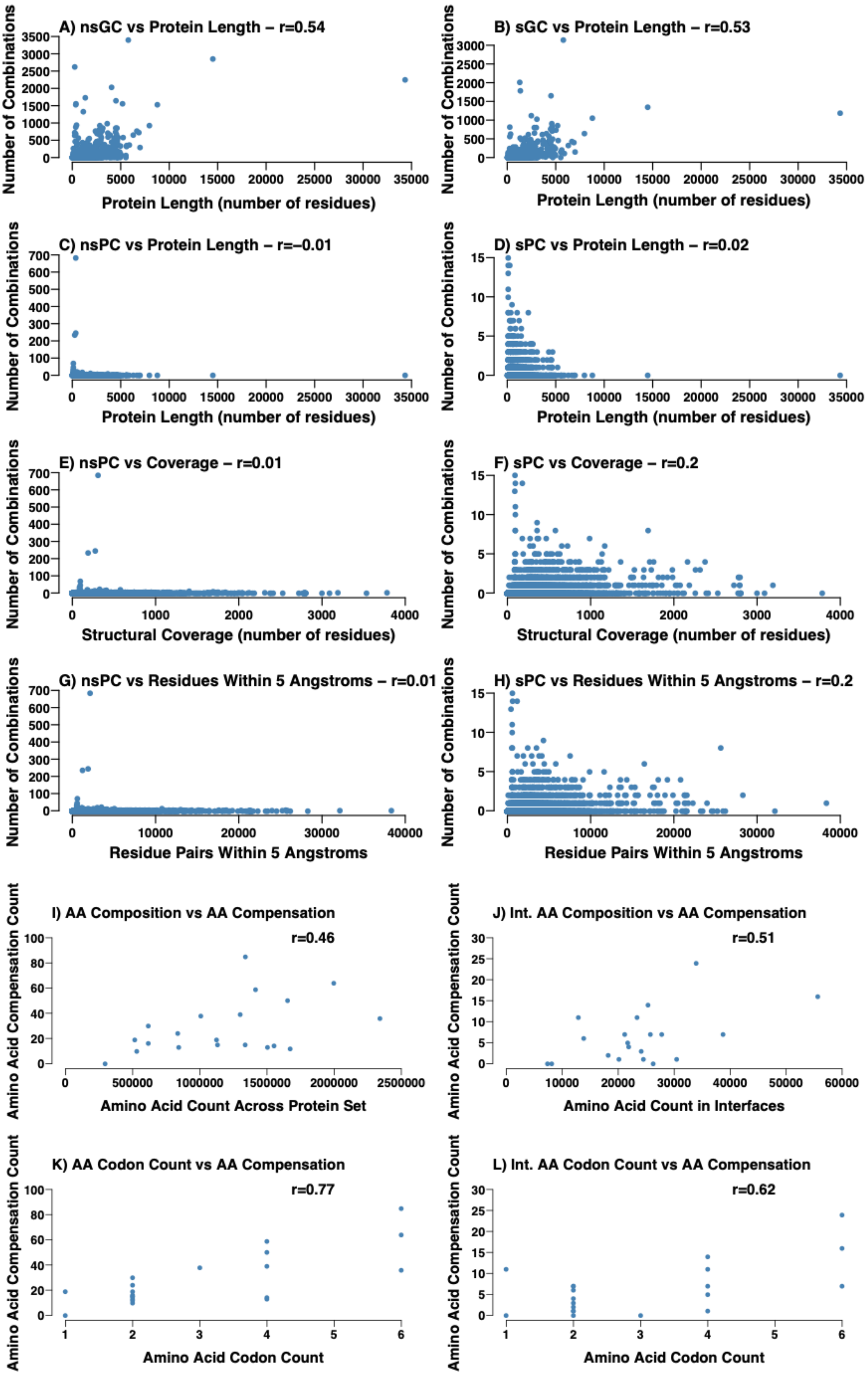
A) nsGC vs Protein Length – r=0.54
B) sGC vs Protein Length – r=0.53
C) nsPC vs Protein Length – r=–0.01
D) sPC vs Protein Length – r=0.02
E) nsPC vs Coverage – r=0.01
F) sPC vs Coverage – r=0.2
G) nsPC vs Residues Within 5 Angstroms – r=0.01
H) sPC vs Residues Within 5 Angstroms – r=0.2
I) AA Composition vs AA Compensation
r=0.46
J) Int. AA Composition vs AA Compensation
r=0.51
K) AA Codon Count vs AA Compensation
r=0.77
L) Int. AA Codon Count vs AA Compensation
r=0.62
Number of Combinations
Protein Length (number of residues)
Structural Coverage (number of residues)
Residue Pairs Within 5 Angstroms
Amino Acid Compensation Count
Amino Acid Count Across Protein Set
Amino Acid Count in Interfaces
Amino Acid Codon Count

**Figure S2:** Normalisation of combination numbers per protein and compensation types. **A)** Non-Synonymous Global Combination number vs protein length. **B)** Synonymous Global Combination number vs protein length. **C)** Non-Synonymous Proximal Combination number vs protein length. **D)** Synonymous Proximal Combination number vs protein length. **E)** Non-Synonymous Proximal Combination number vs structural coverage. **F)** Synonymous Proximal Combination number vs structural coverage. **G)** Non-Synonymous Proximal Combination number vs residue pairs within 5 angstroms. **H)** Synonymous Proximal Combination number vs residue pairs within 5 angstroms. **I)** Numbers of amino acid compensations per amino acid vs numbers of codons per amino acid – Proximal Combinations. **J)** Numbers of amino acid compensations per amino acid vs numbers of codons per amino acid – Interface Variant Combinations. **K)** Numbers of each amino acid across interfaces vs numbers of amino acid compensations per amino acid.

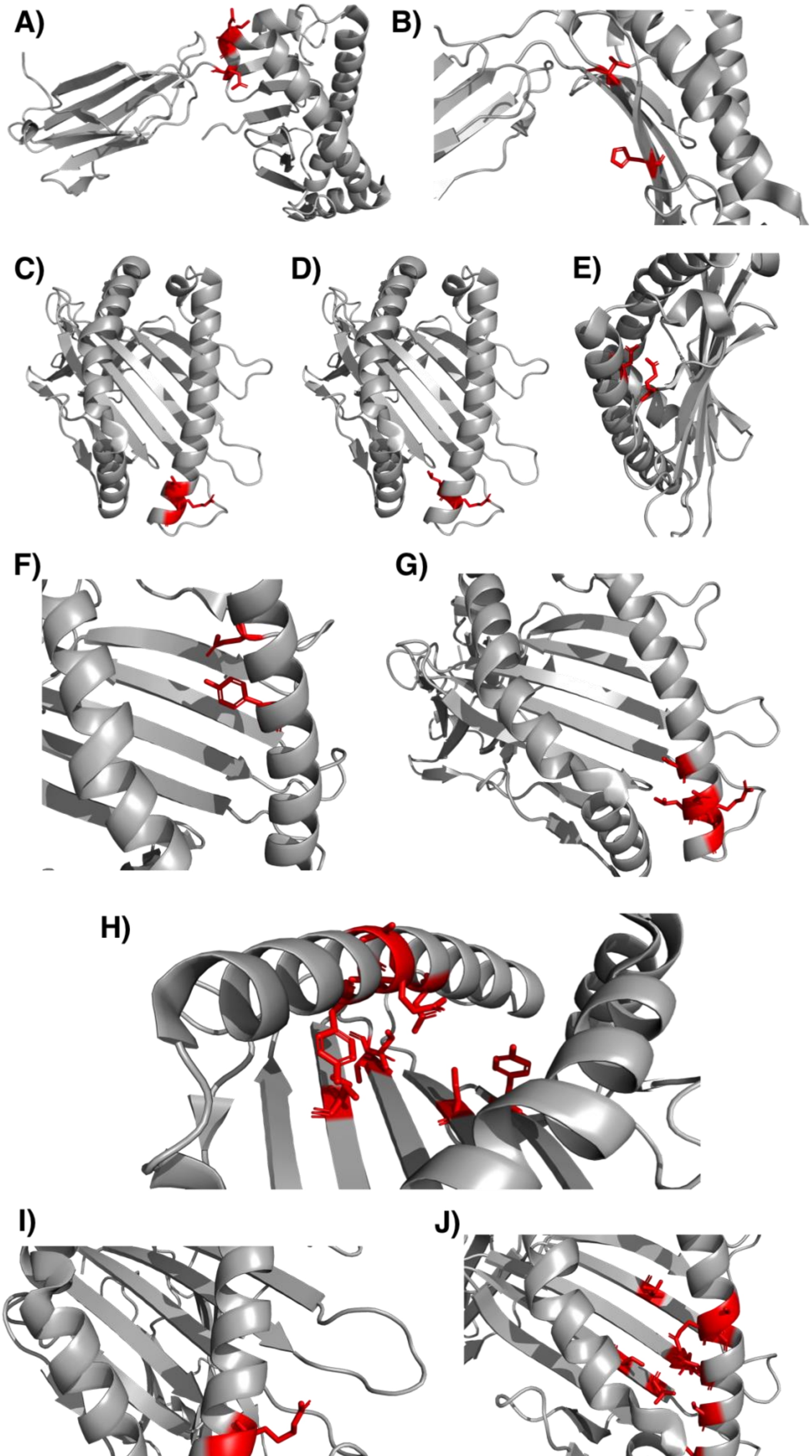
A)
B)
C)
D)
E)
F)
G)
H)
I)
J)

**Figure S3:** Visualisation of the top ten Proximal Combinations of non-synonymous variants in Q31610 by number of total occurrences. Proteins are shown in cartoon format and are coloured grey, and variant positions are shown in stick format and are coloured red. **A)** 'D201E,K202T,E204Q' – 3,972 occurrences. **B)** 'V127L,H137Y' – 416 occurrences. **C)** 'N104I,R106L,G107R' – 393 occurrences. **D)** 'L105A,R106L' – 341 occurrences. **E)** 'E69K,N87E' – 318 occurrences. **F)** 'E69T,Y91F' – 317 occurrences. **G)** 'S101N,N104I,L105A,R106L,G107R' – 278 occurrences. **H)** 'S35A,V36M,S48A,Y91F,A93T,Q94N,A95T,D98Y,S121R,Y140F' – 176 occurrences. **I)** 'R106L,G107R' – 157 occurrences. **J)** 'S35A,V36M,S48A,A93T,Q94N,A95T,D98Y,S101N,L105A, L119W,S121T' – 154 occurrences.

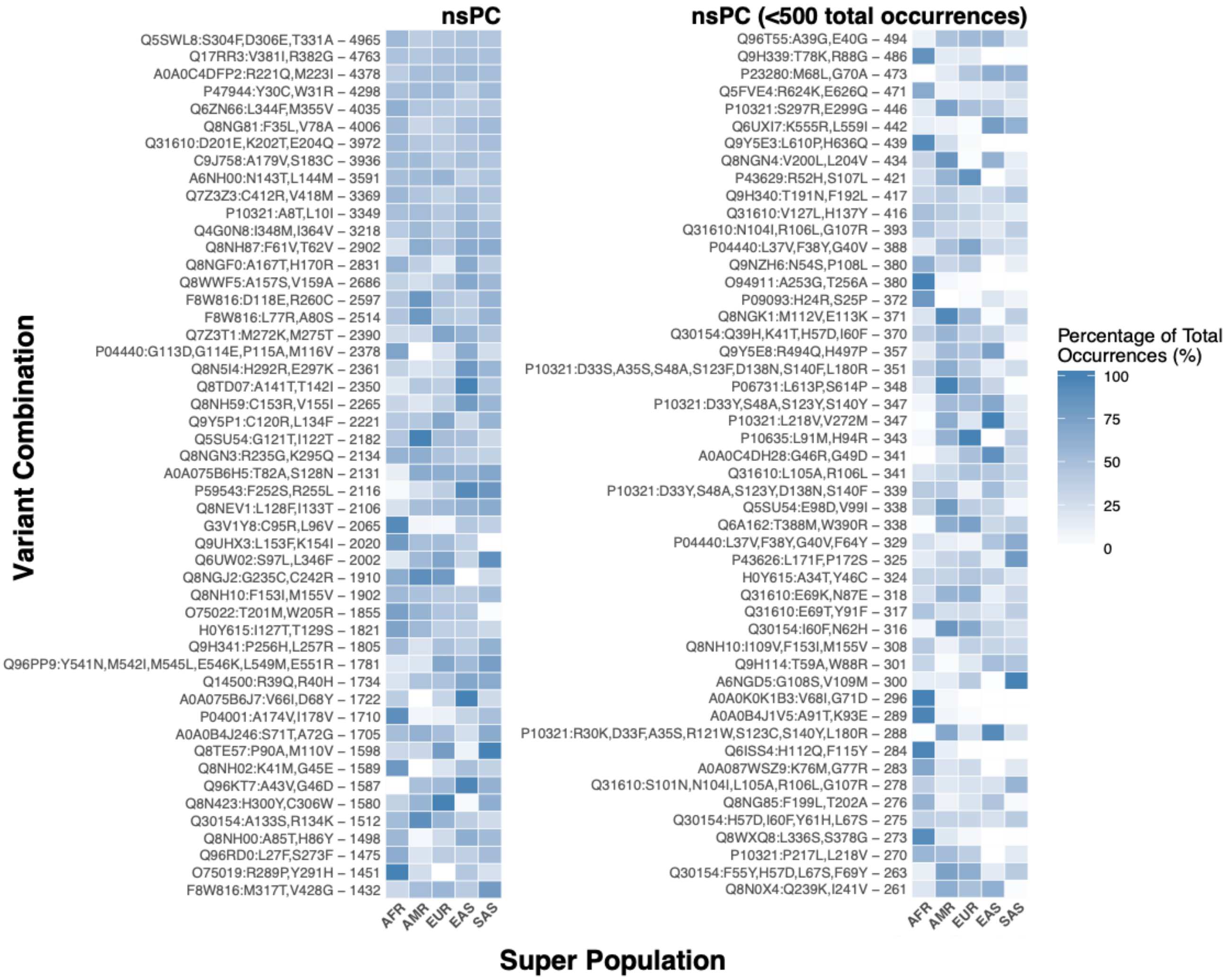


**Figure S4:** Heatmap of occurrences within super populations of the 50 most common Non-Synonymous Proximal Combinations and the 50 most common Non-Synonymous Proximal Combinations for combinations with <500 total occurrences. Combinations are given on the y-axis, see Methods for a description of the variant combination notation format. The number given after the variant combination is the total number of occurrences of the combination for the whole sample set. AFR – African super population, AMR – American super population, EAS – East Asian super population, EUR – European super population, SAS – South Asian super population.

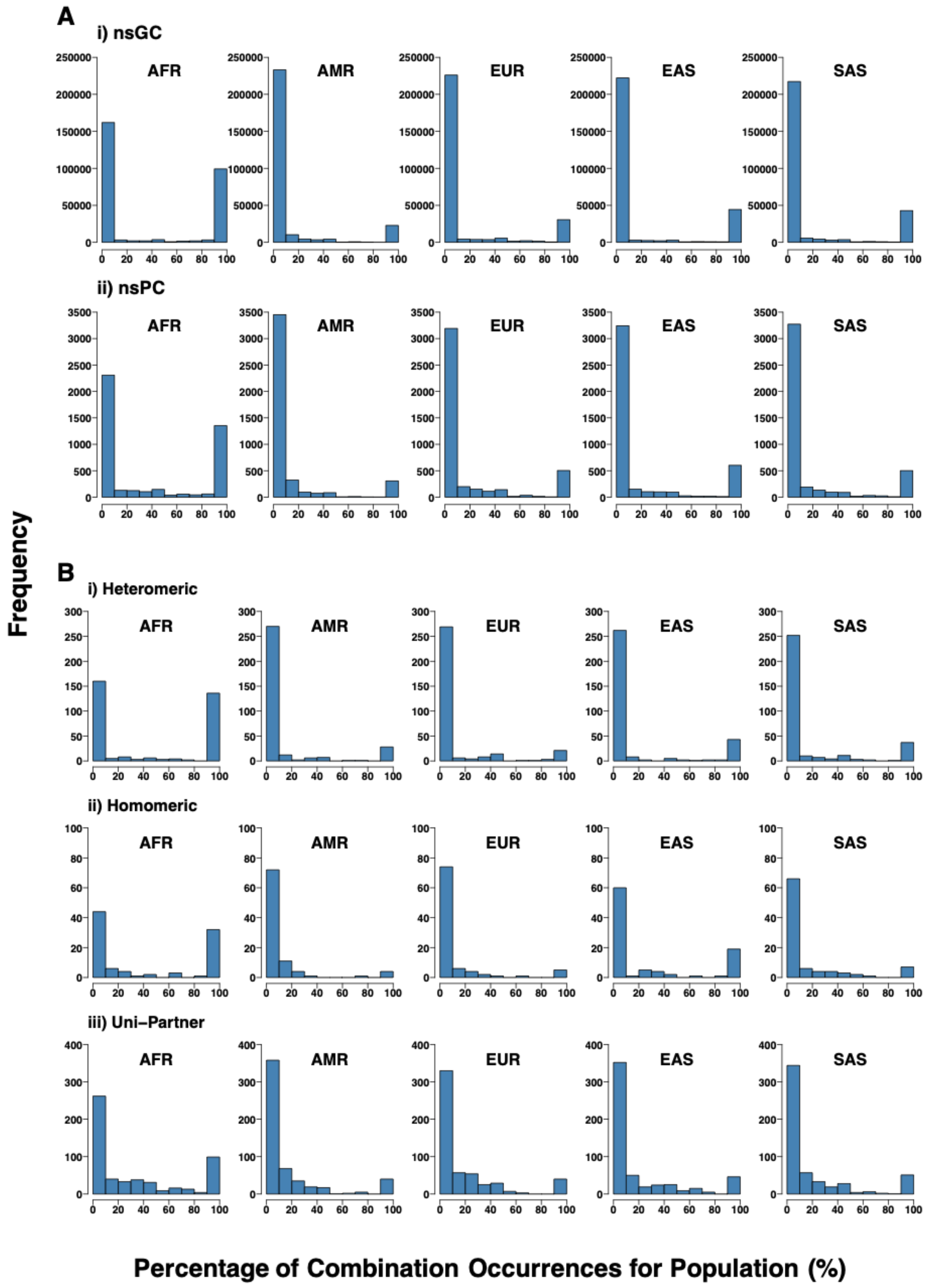


**Figure S5:** Distributions of percentages of total occurrences of each unique variant combination from each super population. **A)** Proximal Combinations. i) Non-Synonymous Global Combinations. ii) Non-Synonymous Proximal Combinations. **B)** Interface Variant Combinations. i) Non-Synonymous Heteromeric. ii) Non-Synonymous Homomeric

Combinations. ii) Non-Synonymous Uni-Partner Combinations. AFR – African super population, AMR – American super population, EAS – East Asian super population, EUR – European super population, SAS – South Asian super population. Note – y-axes differ between the subplots.

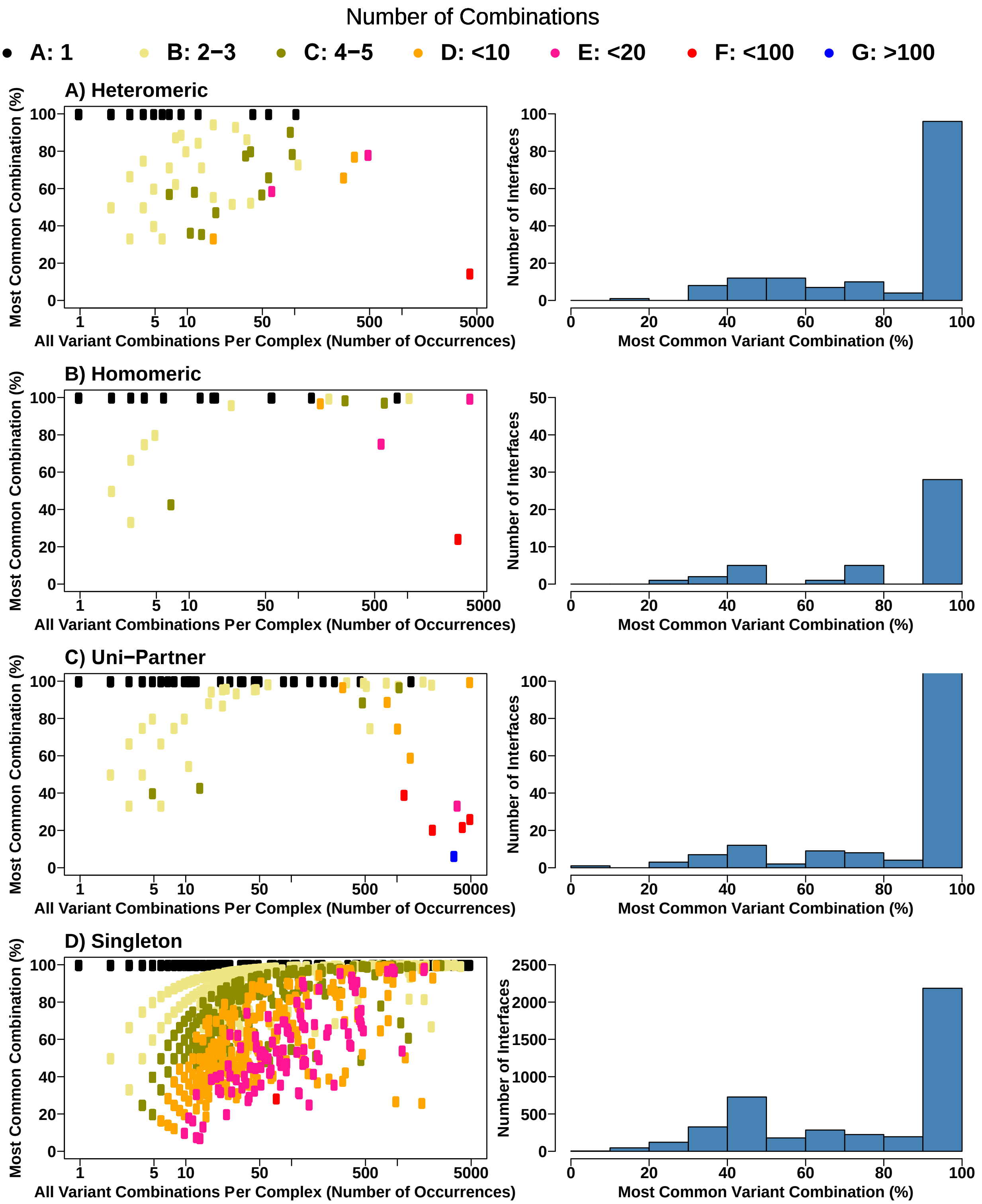


**Figure S6:** Occurrences of the most common non-synonymous interface variant combinations vs all interface variant combinations for each complex. For sub-plots in the left column, each point represents a protein-protein complex, and the colour of the point is determined by the number of unique variant combinations observed for the complex, see legend. The x-axis position of the point is determined by the number of occurrences of any variant combination within the complex, and the y-axis position is determined by the percentage of total combination occurrences accounted for by the most common variant combination in the complex. If the most common variant combination accounts for all of the variant combinations in the complex (black points) the point will lie at the very top of the y-axis (100% of combination occurrences are the most common combination). Sub-plots in the right column show the distributions of proportions of interface variant combination occurrences from the most common variant combination per complex. **A)** Non-Synonymous Heteromeric combinations. **B)** Non-Synonymous Homomeric combinations. **C)** Non-Synonymous Uni-Partner combinations. **D)** Non-Synonymous Singletons. Note – x- and y-axes differ in their scales between the subplots.

### Supplementary Tables

**Table S3:** Properties of amino acids used for finding compensatory amino acid changes

| Amino Acid | Charge | Functional Group | Average Atomic Mass |
|---|---|---|---|
| A | Hydrophobic | Hydrophobic | 71.08 |
| C | Hydrophobic | Hydrophobic | 103.14 |
| D | Negative | Carboxylate | 115.09 |
| E | Negative | Carboxylate | 129.12 |
| F | Hydrophobic | Phenyl | 147.18 |
| G | Hydrophobic | Hydrophobic | 57.05 |
| H | Positive | Positive | 137.14 |
| I | Hydrophobic | Hydrophobic | 113.16 |
| K | Positive | Positive | 128.17 |
| L | Hydrophobic | Hydrophobic | 113.16 |
| M | Hydrophobic | Hydrophobic | 131.19 |
| N | Negative Polar | Amido | 114.10 |
| P | Hydrophobic | Hydrophobic | 97.12 |
| Q | Negative Polar | Amido | 128.13 |
| R | Positive | Positive | 156.19 |
| S | Positive Polar | Hydroxyl | 87.08 |
| T | Positive Polar | Hydroxyl | 101.11 |
| V | Hydrophobic | Hydrophobic | 99.13 |
| W | Hydrophobic | Phenyl | 186.21 |
| Y | Hydrophobic | Hydroxyl & Phenyl | 163.18 |
